# Novel Insights into Post-Myocardial Infarction Cardiac Remodeling through Algorithmic Detection of Cell-Type Composition Shifts

**DOI:** 10.1101/2024.08.09.607400

**Authors:** Brian Gural, Logan Kirkland, Abbey Hockett, Peyton Sandroni, Jiandong Zhang, Manuel Rosa-Garrido, Samantha K. Swift, Douglas Chapski, Michael A Flinn, Caitlin C O’Meara, Thomas M Vondriska, Michaela Patterson, Brian C. Jensen, Christoph D Rau

## Abstract

**Background:** Recent advances in single cell sequencing have led to an increased focus on the role of cell-type composition in phenotypic presentation and disease progression. Cell-type composition research in the heart is challenging due to large, frequently multinucleated cardiomyocytes that preclude most single cell approaches from obtaining accurate measurements of cell composition. Our *in silico* studies reveal that ignoring cell type composition when calculating differentially expressed genes (DEGs) can have significant consequences. For example, a relatively small change in cell abundance of only 10% can result in over 25% of DEGs being false positives.

**Methods:** We have implemented an algorithmic approach that uses snRNAseq datasets as a reference to accurately calculate cell type compositions from bulk RNAseq datasets through robust data cleaning, gene selection, and multi-sample cross-subject and cross-cell-type deconvolution. We applied our approach to cardiomyocyte-specific α1A adrenergic receptor (CM-α1A-AR) knockout mice. 8-12 week-old mice (either WT or CM-α1A-KO) were subjected to permanent left coronary artery (LCA) ligation or sham surgery (n=4 per group). Transcriptomes from the infarct border zones were collected 3 days later and analyzed using our algorithm to determine cell-type abundances, corrected differential expression calculations using DESeq2, and validated these findings using RNAscope.

**Results:** Uncorrected DEGs for the CM-α1A-KO X LCA interaction term featured many cell-type specific genes such as *Timp4* (fibroblasts) and *Aplnr* (cardiomyocytes) and overall GO enrichment for terms pertaining to cardiomyocyte differentiation (P=3.1E-4). Using our algorithm, we observe a striking loss of cardiomyocytes and gain in fibroblasts in the α1A-KO + LCA mice that was not recapitulated in WT + LCA animals, although we did observe a similar increase in macrophage abundance in both conditions. This recapitulates prior results that showed a much more severe heart failure phenotype in CM-α1A-KO + LCA mice. Following correction for cell-type, our DEGs now highlight a novel set of genes enriched for GO terms such as cardiac contraction (P=3.7E-5) and actin filament organization (P=6.3E-5).

**Conclusions:** Our algorithm identifies and corrects for cell-type abundance in bulk RNAseq datasets opening new avenues for research on novel genes and pathways as well as an improved understanding of the role of cardiac cell types in cardiovascular disease.

## Introduction

The endogenous catecholamines epinephrine and norepinephrine activate two classes of adrenergic receptors (ARs) in the heart, β-ARs and α1-ARs. Chronic hyperstimulation of cardiomyocyte β1-ARs contributes to the pathobiology of heart failure^1^. In contrast, we and others have shown that α1-AR activation protects cardiomyocytes against multiple forms of injury both *in vitro* and *in vivo*^2,3^. α1-ARs exist as three molecular subtypes: α1A, α1B and α1D. Each subtype is activated by epinephrine and norepinephrine but exhibits distinct tissue localization and cellular signaling. Within the rodent and human heart, the α1A and α1B subtypes are expressed on cardiomyocytes^4,5^ whereas the α1D-AR subtype is largely found in coronary artery smooth muscle^6,7^.

A burgeoning body of evidence indicates that the α1A subtype mediates the cardioprotective effects of non-selective α1-AR activation^8^. To test whether these adaptive effects require activation of cardiomyocyte α1As, we recently created a cardiomyocyte-specific α1A-knockout mouse line (*Myh6-Cre*x*Adra1a*^fl/fl^ or cmAKO). The cmAKO mice have no discernible basal phenotype but exhibit early mortality and exaggerated pathological ventricular remodeling after myocardial infarction (MI) driven at least in part by unrestrained necroptosis in cmAKO hearts^9^. Unpublished bulk RNAseq suggested intriguing transcriptomic differences between wild-type (WT) and cmAKO mice. However, definitive interpretation of these data is complicated by the fact that MI induces numerous significant changes in the cellular composition of the post-infarct heart including contemporaneous attrition of some cell types and proliferation or infiltration of others. These dramatic changes yield a cellular landscape that is highly heterogeneous, making it difficult to discern whether an observed change in gene expression in bulk tissue is due to changes in the proportional abundance of the cell type(s) that express the gene, altered regulation of the gene itself, or, most likely, a combination of the two. In this setting, where both cell-specific gene expression and cellular abundance are changing, it is a substantial challenge to even ascertain which cell type(s) are expressing a differentially expressed gene and/or driving pathway activation, significantly limiting potential inferences about underlying mechanisms ^10–12^.

Other studies have frequently used single-cell and -nucleus RNA sequencing to evaluate heterogeneous tissues, but these approaches have limitations in the heart due to the size and variable nuclearity of the cardiomyocyte. Cardiomyocytes are over 100 µm in length^13^, precluding scRNA seq approaches as they will not fit through the narrow nozzles of commonly used microfluidics-based approaches such as 10x Genomics^14^. At the same time cardiac cells are variably multinucleated as a product of genetic and environmental factors^15^ (e.g. a myocardial infarction^16^), limiting the use of snRNA seq approaches as it is currently impossible to accurately link observed nuclei count to cell counts in addition to the complications of comparing nuclear-level to cellular-level sequencing data. Beyond these cardiomyocyte-specific issues, we are also aware of studies which report that single cell- and nucleus-RNA sequencing approaches often do not obtain representative samplings of the tissue in question due to variable efficacy in cell isolation between different cell types when constructing a single-cell suspension^17,18^. In contrast, bulk RNA sequencing preserves the cellular makeup of whole tissue but masks the specific contributions of each cell type to the overall expression profile^19^. Brought together, however, the strengths of each approach (cell-type specific expression in single nucleus RNAseq and intact global transcriptomic measures from bulk RNAseq) can be leveraged in reference-based deconvolution while to minimize their respective weaknesses^20^.

In this article, we describe a novel method of computationally estimating cellular proportions from bulk RNA sequencing of cardiac tissue. Our approach uses cell-type-specific expression markers to infer the cellular makeup underlying bulk expression from heterogeneous tissue samples, offering a novel means of parsing out the transcriptomic and cellular response of remodeled cardiac tissue. We first identify highly specific cell-type gene expression markers from a snRNAseq reference panel. Then, we validate our analysis pipeline by applying those markers in the deconvolution of pure cell-type ground truth datasets, before estimating cellular abundances from bulk RNAseq from the left ventricles of WT and cmAKO mice, after either left coronary artery (LCA) ligation or sham surgery. We find more exaggerated changes in cardiomyocyte, fibroblast, and immune cell populations in the cmAKO cohort, consistent with a broadly cardioprotective effect of cardiomyocyte α1A-ARs. We simulate the effect of including cell type proportions in differential gene expression analysis of compositionally distinct bulk RNAseq replicates then apply this method of accounting for cellular composition to characterize the transcriptomic changes between our genotype and treatment groups. We find that many expression changes originally attributed to treatment or genotype are better explained by shifts in cellular proportions. Finally, we use RNAscope with IHC to experimentally validate several genes whose significance was altered after adjustment, confirming that several of the most significantly observed transcriptomic changes are accompanied by cell-type-of-origin proliferation or loss. Requiring only pre-existing data types and no additional experimentation, our method offers a convenient, flexible, and affordable approach to simultaneously characterize the cellular state and abundance changes between compositionally distinct groups and has broad applications beyond cardiac data.

## Materials and Methods

### Mouse husbandry and model generation

cmAKO mice: C57BL/6J and ROSAmT/mG (stock # 007676) mice were purchased from Jackson lab. The cmAKO mouse line was generated by breeding Myh6-Cre (originally from the lab of E. Dale Abel at the University of Iowa and provided by Leslie Leinwand at the University of Colorado) to Adra1a flox/flox mice with loxP sites flanking the first coding exon (constructed in the Paul C. Simpson lab at the University of California, San Francisco/San Francisco VA Medical Center)^21,22^. All mice were backcrossed regularly and maintained on a C57BL/6 genetic background. Twelve-to 16-week-old males were used to generate the myocardial infarction and sham-operated mouse models. Floxed mice (αMHC-Creneg/α1Afl/fl) were used as wild-type (WT) controls for cmAKO mice.

snRNAseq experiments: C57BL/6J mice (JAX stock 000664) were purchased from Jackson Laboratory, Bar Harbor, Maine, and bred to produce pups.

Pure Cell fractions: For experiments involving pure cell-type fractions, C57BL/6J mice were obtained from Jackson Laboratories at 7 weeks of age and allowed to acclimatize for at least two weeks.

### Myocardial infarction model

Mice were subjected to permanent LCA ligation as previously described^9^. In brief, to induce myocardial infarction (MI), a left thoracotomy was performed at the fourth-fifth intercostal space, followed by permanent ligation of the left anterior descending (LAD) coronary using a 7/0 non-absorbable ethylene suture. Occlusion was verified by anemia and akinesis of the apex and anterior-lateral wall, after which the thorax was closed in layers. After extubation, mice were kept warm until fully recovered. For LCA ligation, sham surgery, and terminal cardiectomy, mice were anesthetized by inhalation of isoflurane (2%). For postoperative analgesia, 5 mg meloxicam/kg body weight was applied every 24 hours for the first 72 hours post-surgery.

### Tissue Collection and Preparation

After sacrifice, the heart was quickly excised and sectioned perpendicular to the long axis of the left ventricle. The infarct border zone was visualized under a dissecting microscope and dissected out then immediately snap frozen in liquid nitrogen. Tissue was taken from an anatomically analogous location in sham-operated hearts using an identical process. RNA was extracted and shipped to Novogene (Sacramento, CA) for bulk RNAseq as described below.

To fix heart tissue for immunohistochemistry, mice were heparinized and the heart was perfused with 10 mL of PBS followed by 20 mL of 4% paraformaldehyde (PFA)–PBS through a 23-gauge butterfly needle, then excised and placed in 4% PFA-PBS for 24 hours before transfer to 70% ethanol. Hearts were then embedded in paraffin and 5 µm sections collected.

### Fluorescent In*-Situ* Hybridization (FISH)

Formalin-fixed paraffin sections were used for localizing the cellular expression of various mRNA transcripts by *in situ* hybridization with the RNAscope® Assay^23^ as described by the manufacturer (Advanced Cell Diagnostics, Inc., Newark, CA). Briefly, slides were baked at 60°C for 1 h, deparaffinized with xylene and absolute ethanol, and pretreated with Target Retrieval Reagent, H_2_O_2_, and Protease Plus according to manufacturer specified conditions and times. This was followed by hybridization with RNAscope Probes for 2 h at 40°C in the HybEZ oven for detection of mRNA, or a negative control probe (**Supp. Info.**). RNAscope Multiplex Fluorescent Reagent Kit v2 (Advanced Cell Diagnostics, Inc., Newark, CA, Cat No. 323120) was employed for signal amplification and detection. This kit uses fluorescent probes for the development of a reaction product visible at the Cy5 and Cy7 channels. FITC was not used due to green channel autofluorescence common in the heart.

### Immunofluorescence Staining and Image Acquisition

Immediately following FISH, slides were blocked with 10% normal goat serum for 1 h and incubated with primary antibody overnight at 4°C (**Supp. Info.**) Next, slides were incubated with secondary antibody for 2 hr at RT. Slides were then rinsed in PBS and mounting medium with DAPI was applied and coverslips were attached and sealed with clear nail polish. FISH and IF staining was performed in the University of North Carolina Histology Research Core.

Stained sections were imaged with an Olympus VS200 slide scanner equipped with a motorized stage, an Olympus DP74 digital camera, and OlyVIA software (Olympus America Inc., Center Valley, PA, RRID: SCR_016167). These full scans were opened using Qupath (RRID: SCR_018257), and single field images were generated^24^. 40X full scans are available as supplementary figures.

### Cardiac Cell Isolation for Pure Cell Type Fractions

As previously described, adult B6 mice were treated with heparin (100 USP units) for 20 minutes to inhibit blood coagulation followed by anesthesia with sodium pentobarbital (100 µL of 50 mg/ml dilution, intraperitoneal)^25^. Upon loss of rear foot reflex, hearts were removed and immediately submerged in ice-cold PBS to arrest the heart.

#### Cardiomyocytes

Hearts were mounted on a modified Langendorff apparatus and perfused for 5 minutes with Tyrode’s solution (10u mM NaCl, 5.4 mM KCl, 1 mM MgCl2, 0.6 mM Na2HPO4, 10 mM glucose, 10 mM HEPES (pH 7.37), oxygenated with 95% (v/v) O2 and 5% (v/v) CO2) at 37°C, then perfused for 15-30 minutes with 30 mL Tyrode’s containing 20 mg collagenase type-II and 3mg protease type-XIV, then washed for an additional ten minutes with Krebs buffer (25 mM KCl, 10 mM KH_2_PO_4_, 2 mM MgSO_4_, 20 mM glucose, 20 mM taurine, 5 mM creatinine, 100 mM potassium glutamate, 10 mM aspartic acid, 0.5 mM EGTA, 5 mM HEPES (pH 7.18) oxygenated with 95% O2 and 5 % CO2 (v/v)). Cardiomyocytes were then dissociated in Krebs buffer, filtered through a 100 µm strainer, and centrifuged 2 minutes at 1000G before being placed in triZoL for RNA isolation.

#### Fibroblasts

Fibroblasts were isolated through enzymatic digestion using Liberase (Roche, 5401119001). Hearts were minced into small pieces and transferred into 18 mL of 1x Liberase in Hanks (+Ca, +Mg) media. Solution is gently stirred while incubating at 37°C for 3 minutes. Tissue is allowed to settle, then the supernatant is sieved through a 70 µm strainer and transferred to a tube containing 10 mL of Krebs-Henseleit (KH) buffer (Sigma, K3753). Cell suspension is centrifuged at 1000 rpm for 5 minutes in a (Sorvall Legend Micro 17 Centrifuge). Supernatant was removed and pellet washed with 10 mL ice-cold KH buffer, followed by another round of centrifugation and resuspension in 5 mL of cold KH buffer. The process was repeated using the rest of the heart tissue 4 times until no clumps of heart tissue remained in the original tube. All cells were centrifuged then plated on untreated plates in DMEM/F12 media supplemented with 10% FBS, 1% Pen/Strep and 0.1% insulin-transferrin-selenium (ITS; Corning, 354350). After 2 h, human basic fibroblast growth factor (1:10,000 concentration from a 200x stock, MilliporeSigma, 11123149001) was added to the media. Media was removed after 24 h, plates were washed with PBS, then triZoL was added for RNA isolation.

#### Endothelial Cells

Hearts were prepared as described for fibroblasts, however before plating, precoated CD31 antibody magnetic beads (Miltenyi Biotec, 130-097-418) were introduced to the suspension and incubated for 20 minutes before being immobilized by a magnet and the other cells washed away. Remaining cells were released from the beads, cultured on treated plates with EBM-2 cell media (w/ 10% FBS, 1% pen/strep, 0.1% ITS) for 24 h, followed by wash with PBS to remove dead cells and residual beads, followed by addition of triZoL for RNA isolation. *RNA Isolation*: All RNA isolations were performed using the Zymogenetics Direct-zol RNA miniprep kit (R2052) according to manufacturer instructions. RNA quantity was measured using Qubit RNA High Sensitivity Assay (ThermoFisher, Q32855) and integrity determined using an Agilent Bioanalyzer. Only samples with RIN > 7.0 were used for library preparation, detailed below.

### Cardiac Cell Isolation for snRNAseq

On two occasions, hearts were excised from 3 littermates collected at P21. If 2 females and 1 male were used for the first collection, then the reverse was done on the second collection, such that the final sequencing represents 6 hearts, 3 males and 3 females. Excised hearts, with atria removed, were Langendorff perfused with 25 mLs of 1 mg/mL collagenase as described below.

Hearts were extracted from euthanized mice by cutting the aorta just below the arch arteries, along with the other major vessels. Isolated hearts were washed in ice cold Kruftbruhe (KB) solution and secured by their aortas to a cannula of varying sizes (see Table 2) then tied off with a 3-0 silk suture. Atria were removed with Vannas micro spring scissors. Cannulated ventricles were then hung from a Langendorff apparatus and perfused with calcium-free Tyrode’s buffer, followed by 1 mg/mL collagenase type II (Thermo Fisher, 17101015) dissolved in calcium-free Tyrode’s buffer. Both solutions were warmed to 37°C. Volume of collagenase solution, along with size of cannula, varied by age of mouse (see Table 2). Following perfusion, ventricular tissue was diced with dissection scissors, triturated in ice cold Kruftbrühe (KB) solution using a wide bore 1 mL pipette

Digested hearts were resuspended in ice cold KB and allowed to settle for 10 minutes on ice. The supernatant was removed and the loose pellet was resuspended in 5 mL of Lysis buffer prepared as previously described^26^, with only one adjustment – 50 µl of 10% Triton-X-100 was added (final concentration 0.1%). Cells were incubated in Lysis buffer + Triton for 5 minutes on ice, after which they were homogenized with a Tissue Tearor electric tissue homogenizer (Model # 985370) at the second lowest setting for 20-30 seconds and left to sit again for another 5 minutes on ice. They were then transferred through a 15 mL glass Dounce homogenizer and further homogenized with 20 strokes of the A pestle and 20 strokes of the B pestle. Homogenized cell suspensions were sequentially filtered through a 70 µM, 40 µm, and 20 µm cell strainer to remove debris and undigested materials. Samples were then spun at 1000G for 5 minutes and resuspended in 1 mL of 2% BSA dissolved in D-PBS with RNaseOut (Invitrogen, 200U/mL). A small aliquot was set aside to serve as an unstained control for fluorescent activated cell sorting (FACS). The remainder of the suspension was stained with DAPI at 10 µg/ml for 5 minutes on ice. Samples were spun at 1000G for 5 minutes and resuspended in fresh 2% BSA-RNaseOut solution.

Following staining, nuclei were sorted on a BD FACSMelody at 4°C. Following standard protocols, forward and side scatters were used to remove doublets. Unstained controls were used to set the V450 gate. 432,000 nuclei were collected into a 2 mL centrifuge tube preloaded with 500 µL of 2% BSA-RNaseOut solution. Sorted nuclei were spun down at 1000G for 5 minutes, supernatant was removed, and samples were resuspended in 100 µL of 2% BSA-RNaseOut solution (Invitrogen) before proceeding to 10x library preparation.

### Bulk and Single Nucleus RNAseq Library Preparation

#### snRNAseq

Nuclei were quantified with a Luna Fl cell counter (Logos Biosystems) and the volume was adjusted to obtain the ideal concentration of nuclei recommended by 10x Genomics (1000 nuclei/µL). Individual nuclei were paired with Chromium v3.1 gel beads and cDNA synthesis, barcoding, and dual index library preparation was performed using Chromium Next GEM V3.1 chemistry according to the manufacturer’s recommendation (10x Genomics). 10,000 nuclei were targeted for each sample with 13 cycles for cDNA amplification and 13 cycles for sample index PCR. The fragment size of cDNA and libraries was assessed using Agilent’s 5200 Fragment Analyzer System to verify product quality prior to sequencing with RIN >7.0 used as a cutoff for sequencing.

#### Bulk RNAseq of purified fractions

Sequencing libraries were prepared using 3 μg total RNA from isolated RNA using the Stranded mRNA-Seq Kit (KAPA Biosystems).

#### Bulk RNAseq of whole tissue

mRNA-Seq libraries were constructed using 4 μg total RNA with the Stranded mRNA-Seq Kit (KAPA Biosystems) at Novogene (Sacramento, CA).

### RNA sequencing

#### snRNAseq

2 libraries (3 mice per library) were sequenced at the Roy J. Carver Biotechnology Center at the University of Illinois, Urbana Champaign on a NovaSeq 6000 using one S4 lane with 2X150nt reads. Samples were demultiplexed and mapped to the mm10 genome using Cell Ranger v6.1.1 (10X Genomics).

#### Bulk RNAseq of purified fractions

Bulk RNA fractions were sequenced 2×50 (paired end) on a Novaseq 6000 S2 chip at the Technology Center for Genomics and Bioinformatics (TCGB) at UCLA.

#### Bulk RNAseq of whole tissue

Samples were multiplexed with Illumina TruSeq adapters and run on a single 75-cycle paired end sequencing run with an Illumina NextSeq-500.

All bulk RNAseq samples were demultiplexed, then decoy-aware pseudo-aligned to the GRCm39 transcriptome with Salmon v1.10.2^27^ and read quality was checked with FastQC v0.12.1.^28^ and summarized with MultiQC^29^. Then, samples were joined into a counts matrix with the txmeta^30^ and tximport^31^ packages in R version 4.3.1. Bulk RNAseq and snRNAseq data are publicly available as NCBI BioProjects under accessions PRJNA1122769 and PRJNA880279 or in SRA SUB14505282 and SRP398524. The full Snakemake^32^ analysis pipeline is publicly available on Github (https://github.com/guralbrian/bulk_decon).

### Single Nucleus RNAseq Data Analysis

Raw counts, barcode, and feature matrices were joined into a single Seurat object^33^ for each replicate. Ambient droplet RNA and doublets were identified and removed *in-silico* with DropletUtils::emptyDrops^34^ and scDblFinder::computeDoubletDensity^35^, respectively. Sample replicates were merged into a single Seurat object, then filtered by feature count, transcript count, mitochondrial transcript percentage, and doublet score. Raw counts were normalized, scaled, applied to principal component analysis, and integrated by harmony::RunHarmony^36^. For clustering, the number of principal components included was determined by finding the first PC which exhibits cumulative percent variability greater than 90% or that which explains less than 5% of the total variability. Samples were then clustered by the standard Seurat methods, using the *harmony* reduction. See Supplementary Data for detailed transcriptional marker list.

### Deconvolution analysis

We converted Ensembl IDs to gene symbols to match the bulk and snRNA-seq formats; among duplicate gene symbols, we kept the one with the highest average gene expression.

Before performing deconvolution, the 35,334 unique transcripts from the bulk RNAseq data and 19,883 from the snRNAseq data were subset to the 15,376 genes present in both datasets. Then, scran::findMarkers^37^ was used to find markers for each Seurat cluster. Markers were ordered by adjusted p-value and the top 15 markers per cluster were retained. After, marker genes were manually queried in ToppGene^38^ for associations with known cell types. High confidence cell type associations were annotated to the relevant cluster and small clusters or those with low confidence annotated were excluded from further analysis. After, five cell-type clusters were retained: endothelial cells, cardiomyocytes, fibroblasts, a joint vascular smooth muscle cell and pericyte cluster, a joint monocyte and macrophage cluster, and smooth muscle cells.

Deconvolution of all bulk RNAseq datasets was performed with MuSiC^39^, an R package for deconvolution of bulk RNA sequencing data which uses single cell or single nucleus RNA sequencing data as a reference. Only genes present in the 15 markers per cell type identified with scran (**Supplemental Data 1**) were included in deconvolution analysis. Raw expression counts were applied to MuSiC::music_prop and proportion estimates from the weighted deconvolution were used in all further analysis.

### Dirichlet Model

To model the relationship between treatment or genotype with estimated cell type proportions, we used a Dirichlet regression model. The DirichReg function from the DirichletReg^40^ package was run with a genotype-treatment interaction term with common parameterization.

### Differential Expression

Differential expression analysis was performed with DESeq2^41^. Two iterations were performed: without covariates, and with covariates including centered log-ratio (clr) transformations of cardiomyocyte and fibroblast proportions. The DESeq2 workflow included creating DESeqDataSet objects, filtering genes with more than 10 reads in at least four samples. Both iterations modeled gene expression as the product of the additive effects of genotype, treatment, and their interaction, with the second analysis iteration also including additive effects of each cell type proportion. Multiple testing correction was performed using a false discovery rate (FDR) threshold of 0.05. See **Supplemental Data 2** for full DESeq2 outputs.

### Simulated Differential Expression

Bulk RNAseq sample groups were generated with predefined cell type contributions to total expression by referencing our snRNAseq dataset. Specifically, a transcriptomic profile was generated for each cell type cluster by summing the expression of all nuclei within the cluster and dividing by the summed transcript counts in that cluster. This yielded probability for each gene being selected if a single transcript was drawn at random from a given cell type cluster. Transcripts were then sampled from these profiles to a total of 25 million reads to mimic typical bulk RNAseq data^42^. 81 groups of four replicates were modeled, with the cardiomyocyte proportion ranging from 30 - 70% cardiomyocytes and the remaining expression evenly sourced from fibroblasts, endothelial cells, macrophages, and pericytes/smooth muscle cells.

To test the effect of including composition, DESeq2 was run with two models:

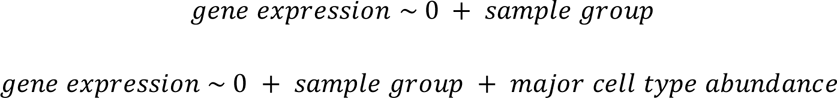

For each model, sample groups were contrasted against the 50% cardiomyocytes group and the number of significantly differentially expressed genes was recorded. The analysis was repeated with a total of three different representations of cell type proportions in the model: untransformed proportions, centered-log-ratios of proportions, and principal components from principal component analysis of cell type proportions. All differential expression analysis were conducted in batches of 500 genes.

### GO Analysis

Expression changes of individual genes were summarized into biological pathways by gene ontology (GO) enrichment using clusterProfiler v4.10.0^43^. Gene names were converted to ENSEMBL IDs using the org.Mm.eg.db reference package v3.18.0^44^. For each model and variable, all genes with an adjusted P-value less than 0.1 and an absolute log-fold-change greater than 0.583 were supplied to clusterProfiler::enrichGO using biological pathway ontology and Benjamini-Hochberg p-value adjustment. See Supplementary Data for entire GO term dataset.

### Figure Generation

Schematics and diagrams (**Figures 1A, 2A, 3A, 3B, 4A, and 4B**) were produced with Biorender. All other plots were generated using the following R packages: dot and bar plots were produced with ggplot2^45^, upset plots with ComplexUpset^46,47^, UMAP plot with ggplot2 and tidyseurat^48^, ternary plot with ggtern^49^, and tables with gt^50^ and gtExtras^51^.

**Figure 1.**
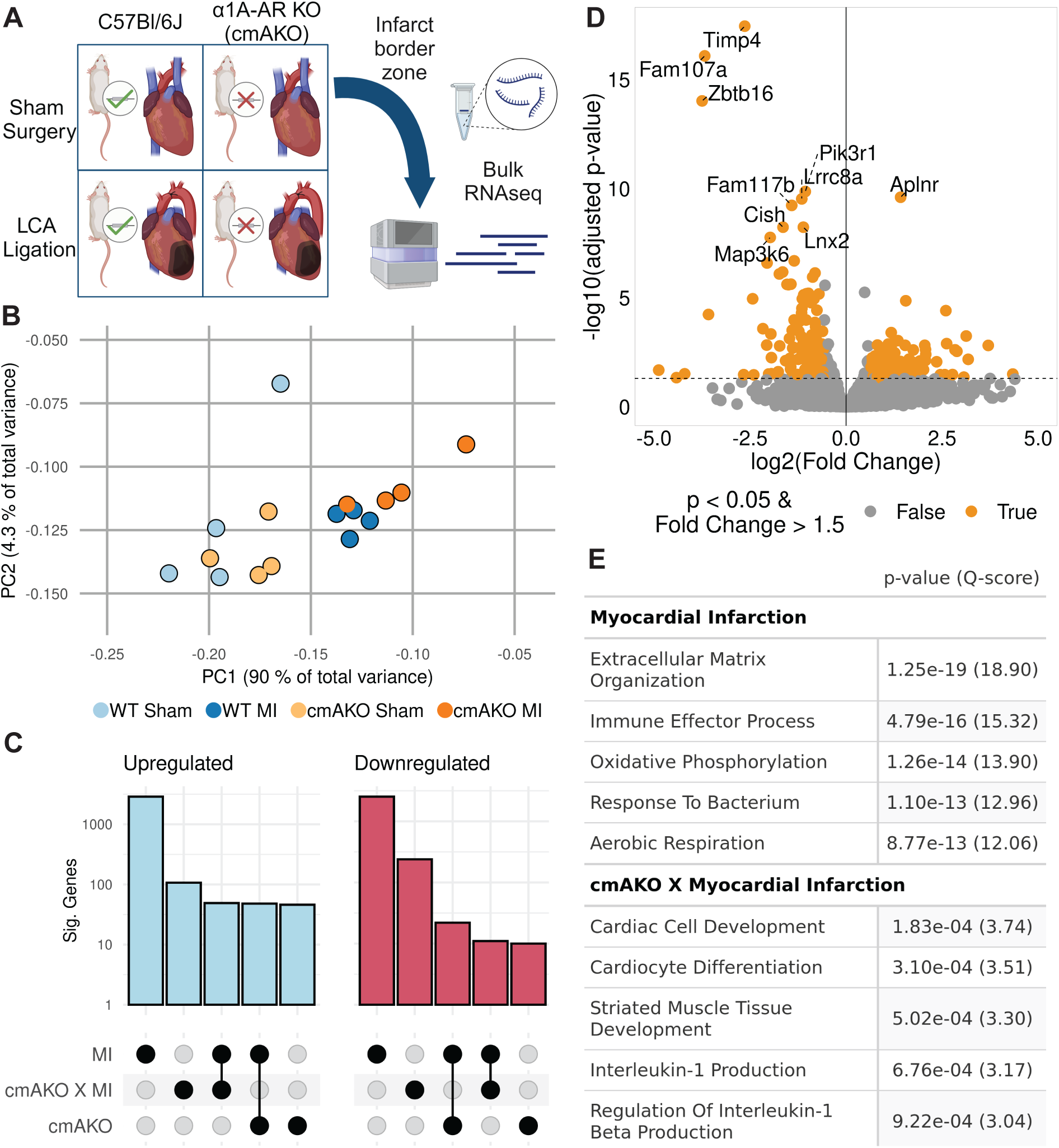
Transcriptomic profiling and analysis in cardiomyocyte-specific α1A-AR knockout mice following myocardial infarction (MI). **A**, Schematic of the experimental design for inducing MI in cmAKO or WT mice and subsequent bulk RNAseq analysis from tissue collected at the infarct border zone and matched sham locations. **B**, Principal Component Analysis (PCA) of RNA counts showing stratification of samples by treatment along PC1, accounting for 90% of the variance. **C**, Differential expression analysis indicating MI as the condition with the highest number of differentially expressed genes (DEGs), followed by the cmAKO and MI interaction; upregulated genes are shown in blue and downregulated genes in red. **D**, Volcano plot of DEGs specific to the cmAKO response to LCA ligation, highlighting genes like Timp4 (fibroblasts) and Aplnr (cardiomyocytes) with p < 0.05 and fold change > 1.5. **E**, Gene Ontology (GO) enrichment analysis showing biological processes significantly influenced by MI, cmAKO, and their interaction, such as extracellular matrix organization and cardiac muscle cell development. Terms with identical p-values have been consolidated.

**Figure 2.**
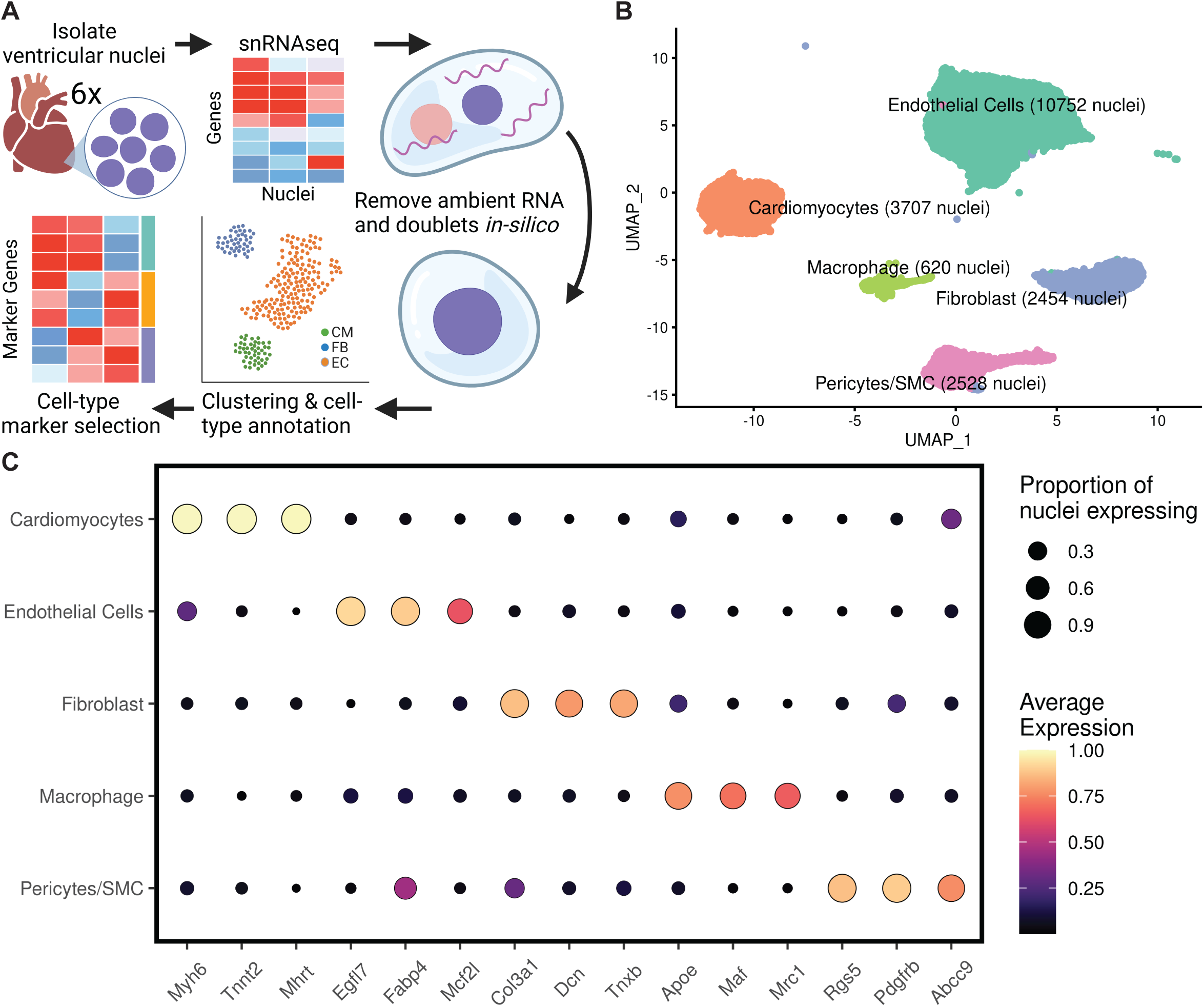
Development and characterization of a cell-type-specific gene expression reference panel from single-nucleus RNA sequencing (snRNAseq). **A,** Generation of the reference panel involved identifying five major cell types in snRNAseq data from pooled nuclei of left ventricles of untreated C57BL/6J mice. After preprocessing, 15 marker genes per cell type were selected based on significance using the findMarkers function from scran. **B,** Annotation and clustering within the snRNAseq reference identified five clusters, each labeled with broad cell type names based on known markers. Post-processing included dimensional reduction and exclusion of clusters with fewer than 400 nuclei or unclear cardiac cell type annotations, resulting in 20,061 nuclei across the five clusters. **C,** Specificity of cell cluster gene expression markers demonstrated by the top three markers for each cell type, selected by log-fold-change in one-versus-all testing between clusters. Marker visualization includes point size proportional to the nuclei expression and color coding by average expression within each cluster.

**Figure 3.**
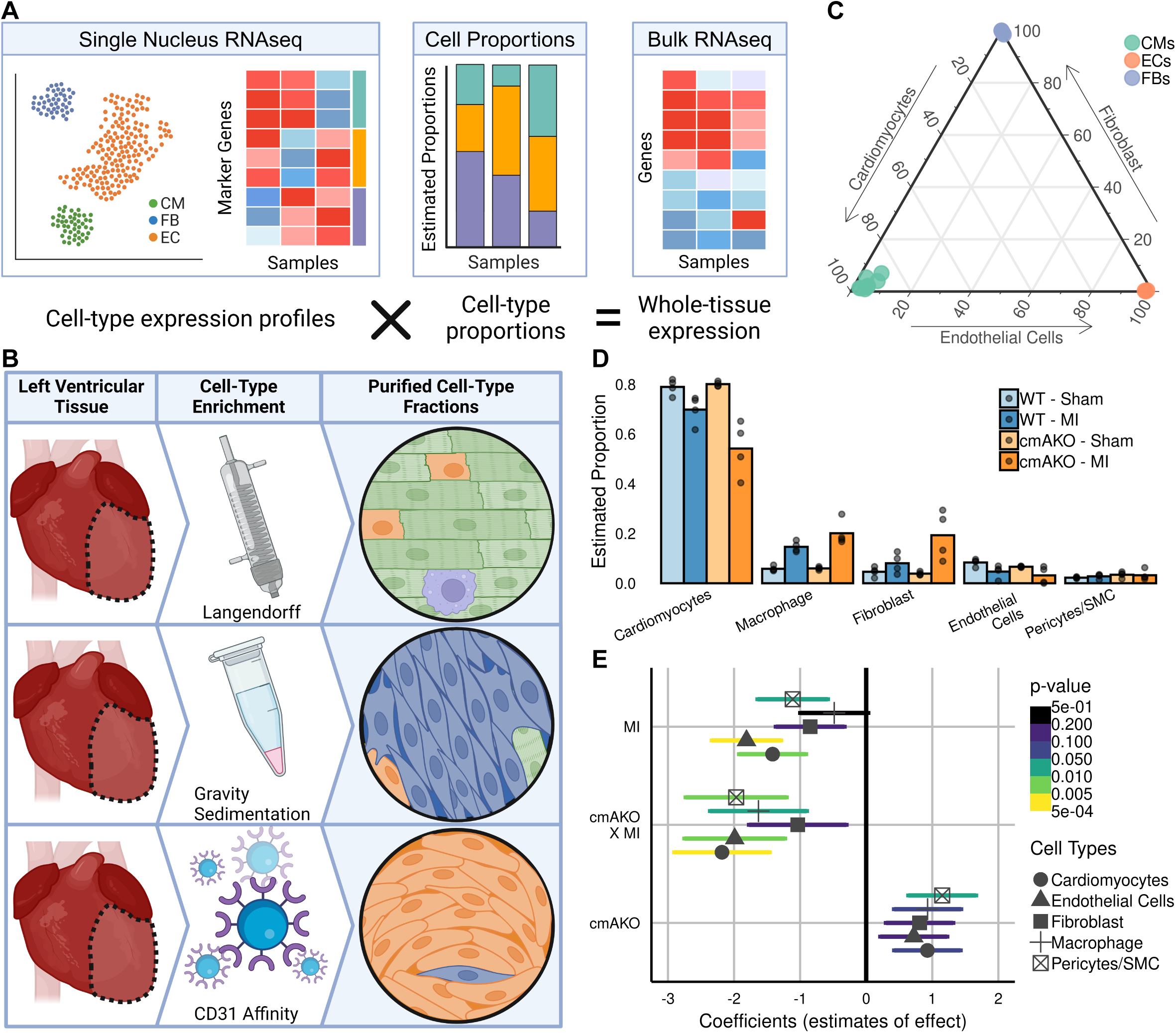
Deconvolution of cell type proportions from bulk RNAseq data and their dynamics during myocardial infarction in cardiomyocyte-specific α1A-AR knockout mice. **A**, Overview of the reference-based deconvolution approach which utilizes gene expression markers to estimate the proportions of cell types contributing to bulk gene expression. **B**, Generation of cell-type enriched bulk RNAseq data involved isolating cardiomyocytes, fibroblasts, and endothelial cells from left ventricles of untreated WT mice using Langendorff perfusion, gravity sedimentation, and CD31-bead binding. **C,** Validation of the deconvolution pipeline using bulk RNAseq from pure cell type fractions as ground-truth samples, depicted in a ternary plot where each dot represents a replicate, color-coded by intended enriched cell type, indicating high accuracy of the composition estimates. **D**, Estimation of cardiac cell type composition changes post-myocardial infarction (MI) and in α1A-AR knockout (cmAKO) mice, showing increased proportions of macrophages and fibroblasts and decreased proportions of cardiomyocytes, with more pronounced changes in cmAKO mice. **E**, Results from a Dirichlet regression modeling cell type proportions as influenced by cmAKO, MI, and their interaction, revealing significant effects on several cardiac cell types. CMs = Cardiomyocytes; ECs = Endothelial Cells; FBs = Fibroblasts; WT = Wild-Type; MI = Myocardial Infarction

**Figure 4.**
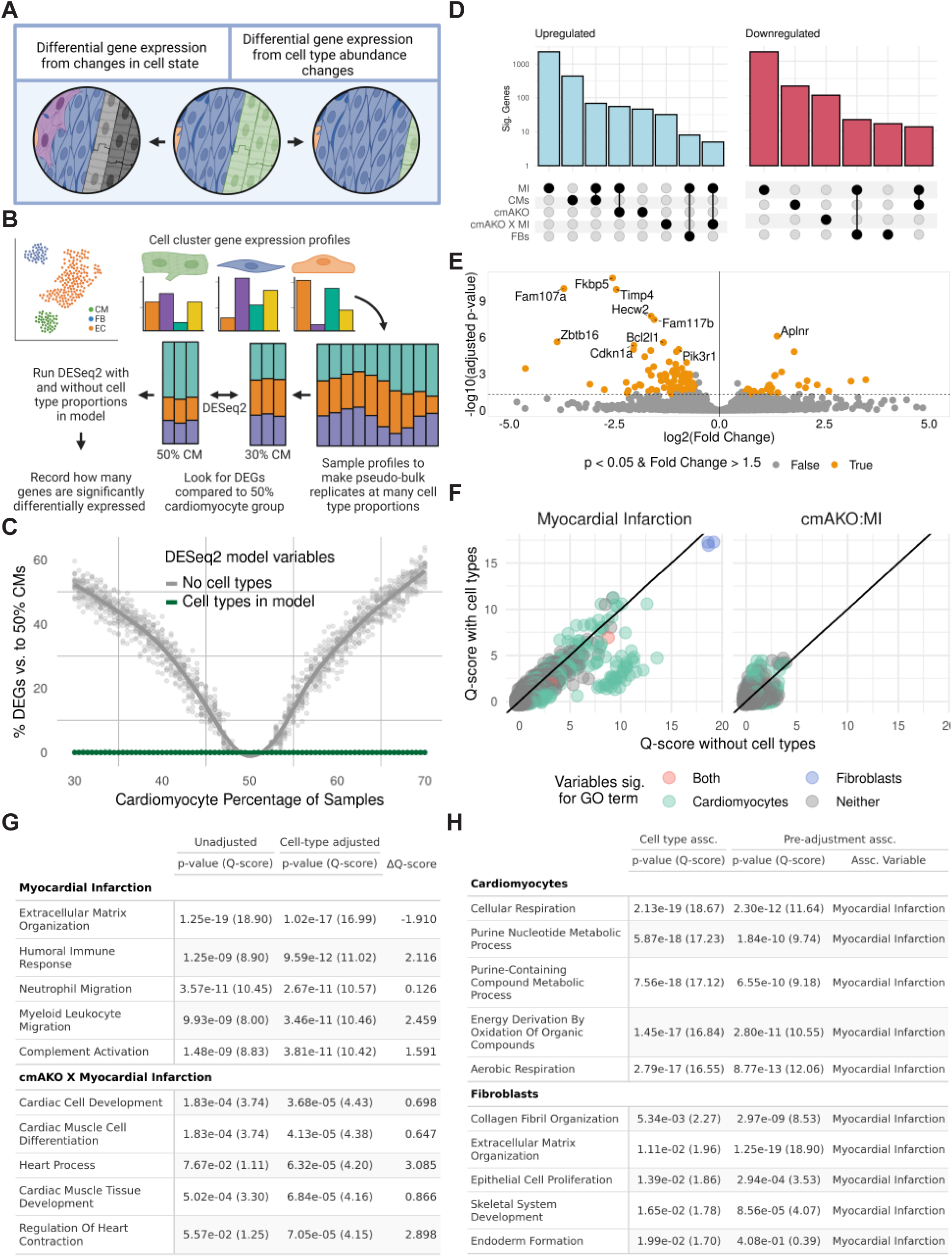
Impact of cell type composition on differential gene expression during myocardial infarction in α1A-AR knockout mice. **A**, Differences in either cell type abundances or novel states can lead to gene expression differences in bulk tissue. B, Schematics of one simulation of compositional changes in differential expression; bulk RNAseq samples were generated with varying cardiomyocyte proportions (30-70%). All samples were compared against the 50% CM group with and without cell-types in the model. **C**, DESeq2 simulations show increases in differentially expressed genes with compositional differences; including cardiomyocyte proportions in the analysis ablates this effect. Each dot is one simulation of 500 genes. **D**, Upset plot showing DEGs related to each variable after adjusting for cell-type composition. Cardiomyocyte proportion is the second strongest asssocaited variable. **E**, Volcano plot showing differential expression of genes associated with cmAKO X MI interaction after adjusting for cell type proportions. **F**, Changes in GO term associations and their significance before and after cell type adjustment, depicted through a scatter plot of Q-scores. Many myocardial infarction-associated GO terms are attributed to changes in cell type abundance. **G** and **H**, Listing of top GO terms post-adjustment showing significant associations with myocardial infarction and cmAKOxMI (**G**), and with changes in cardiomyocytes and fibroblasts (**H**)

## Results

### Unadjusted transcriptional changes in ischemic border zone of cmAKO mice

To examine the role of cardiomyocyte α1A-ARs after heart injury, we created a cardiomyocyte-specific α1A-AR knockout mouse line (*Myh6-Cre*x*Adra1a*^fl/fl^ or cmAKO)^9^. We induced a myocardial infarction (MI) in this model by permanent LCA ligation at 8-12 weeks of age. Three days post-ligation, we collected tissue from the border zone of the infarcted left ventricle, or a matched location in the sham surgery group, and measured bulk gene expression by RNAseq **(Figure 1A)**. To identify outliers and unexpected sources of variation that might bias our results, we performed principal component analysis. Samples were well stratified by treatment across PC1, which accounted for 90% of the variation in gene expression. Although a replicate from the WT sham surgery group deviated on PC2, that axis only represented 4.3% of variation in our data and was included for further analysis (**Figure 1B**). We proceeded with differential expression analysis using DESeq2^41^, modeling gene expression as the sum of the effects of genotype, treatment, and their interaction. LCA ligation produced the greatest number of differentially expressed genes (DEGs), followed by the interaction of cmAKO and LCA ligation, whereas cmAKO alone yielded the fewest DEGs **(Figure 1C>)**. These findings recapitulate our prior studies, where we found that α1A-knockout magnified the adverse effects of LCA ligation but elicited minimal phenotypic differences from WT mice in the absence of cardiac insult^9^.

When examining the genes that are differentially expressed in the cmAKO x myocardial infarction interaction term (the unique response of cmAKO to LCA ligation), we observe a large number of genes that are enriched in specific cell types, such as Timp4 (fibroblasts)^52^, Zbtb16 (cardiomyocytes)^53^, and Aplnr (cardiomyocytes)^54^ (**Figure 1D**). Furthermore, analysis of all DEG with p< 0.1 and FC > 1.5 by gene ontology (GO) enrichment using clusterProfiler^43^ revealed strong enrichments of cardiomyocyte-specific pathways. Likewise, LCA ligation alone showed strong enrichments for cardiac fibrosis and CM energy metabolism (**Figure 1E**). Taken together, these suggest that cell-type compositional changes may contribute to bulk expression changes in the heart. Considering the abundance of expression changes related to cell-type-specific pathways and mechanisms, we next sought to estimate sample-specific changes in cellular composition.

### Cell type-specific markers in snRNAseq

To guide the interpretation of our cmAKO model, we developed a cell-type-specific gene expression reference panel. To this end, we performed single nucleus RNAseq from the left ventricles of WT mice (n = 6). We applied a strict preprocessing and quality control pipeline, which began with *in silico* removal of ambient RNA and low-quality nuclei before proceeding with dimensional reduction, and clustering, retaining 20,968 nuclei assigned to 11 clusters (**Figure 2A, Supp. Fig. 1**). Next, we sought to connect our nuclei clusters to known cell types by comparing cluster-specific expression markers to known cell-type markers. Cluster markers were identified as genes whose within-cluster expression was significantly higher than out-of-cluster expression in one-versus-all testing using findMarkers from scran^35^. The top 15 most significant marker genes per cluster were supplied to ToppGene to look for enrichment of existing cell type markers. After excluding clusters with less than 500 nuclei or those which presented unclear cell-type associations, the final dataset contained 20,061 nuclei assigned to five cell-type clusters. In order of most-to-least abundant, we retained clusters of endothelial cells, cardiomyocytes, fibroblasts, macrophages, and a joint cluster of pericytes and smooth muscle cells (**Figure 2B**). Further, we found that our marker list recapitulates canonical proteomic and transcriptomic cardiac cell-type markers, including Myh6^55^ and Tnnt2^56^ for cardiomyocytes, Egfl7^57^ and Fabp4^58^ for endothelial cells, and Dcn and Col1a1^59^ for fibroblasts (**Figure 2C, Supplemental Data 1**)

### Cell-type composition of mouse left ventricular tissue

Cell-type deconvolution is a method to infer how much RNA comes from each cell type present in bulk RNA sequencing data produced from heterogeneous tissue. For our pipeline, we chose to utilize MuSiC, one of several existing deconvolution methods, which iteratively tests for the combination of cell type proportions whose summed expression profiles best explains the overall tissue expression profile. (**Figure 3A**). To inform development of our analysis pipeline and to test its performance, we first applied it to several bulk RNAseq datasets derived from purified fractions of major cardiac cell types (**Figure 3B**). We found that our MuSiC-based method was able to accurately predict the makeup of every cardiomyocyte, endothelial cell, and fibroblast sample (**Figure 3C**). Confident in our ability to identify major cell types of the heart, we then applied our pipeline to the bulk RNAseq data from our experimental groups. In the untreated WT mice, we estimated that the major cell type *by transcript abundance* was cardiomyocytes (79%), followed by fibroblasts (8.4%) and macrophages (5.8%). For both genotypes, cardiomyocyte abundance decreased in the treatment group, while fibroblasts and macrophages rose in relative abundance. For each of these changes, these compositional shifts were more pronounced in cmAKOs when compared to the WT group (**Figure 3D**).

Traditional models and association tests, such as the Pearson correlation^60^, are prone to identifying spurious correlations when applied to contexts where measures are interrelated, as with cellular composition data^61^. To understand this, imagine a scenario where a treatment causes cardiac fibroblasts to proliferate while having minimal effect on adjacent cell types. If researchers were to quantify the number of each cell type in a treated heart they would find that the absolute number of non-fibroblasts remains constant. However, since the total number of cells in the heart increased, the relative proportion of non-fibroblasts would seem to decrease compared to an untreated heart. As such, standard statistical tests applied to these compositional measures would report that the drug is associated with both an increase in fibroblasts and a decrease in non-fibroblasts, despite there being no absolute increase in the number of non-fibroblasts.

To address this and correctly quantify the effect of each treatment and genotype on the compositional changes, we used a Dirichlet regression model, which can accommodate the sum-to-one constraint and covariance structure of compositional data^40,62^. Our model considered the effects of each genotype and treatment additively, as well as the interaction of the two terms. We found that MI and the cmAKO X MI interaction are the only terms to have significant effects on major cell type proportions. Specifically, MI and cmAKO X MI were both significantly associated with the abundance of cardiomyocytes (p = 0.0051 for MI; p = 0.0024 for cmAKO X MI), endothelial cells (p = 0.00053 for MI; p = 0.00888 for cmAKO X MI), and pericytes/SMC (p = 0.036 for MI; p = 0.01 for cmAKO X MI). Additionally, only cmAKO X MI was found to be significantly associated with the proportion of macrophages (p = 0.026) (**Figure 3E**).

### Cell-type-adjusted Differential Gene Expression

Variation in either cell state or abundance^63^ can be responsible for differences in the overall transcriptional profiles of cellularly heterogeneous tissue (**Figure 4A).** Given the differences in cellular abundance in our samples, we became interested in discerning the contribution of these compositional differences to the transcriptional changes we observed in response to LCA ligation and CM-α1A KO.

To model the inclusion of compositional estimates in gene expression analysis, we simulated bulk RNAseq sample groups composed of a range of underlying cellular proportions (**Figure 4B**). *In silico* mixtures were generated by sampling expression of individual genes from each cell-type cluster according to pre-specified cell-type ratios. We began with 50% cardiomyocytes and equal proportions of each remaining cell type from our snRNAseq panel. We then introduced stepwise changes in cardiomyocyte proportion from a range of 30 - 70% cardiomyocytes, adjusting the other minor cell types as appropriate while adding small amounts of noise to their proportions to avoid full rank modeling errors. We then tested for differential gene expression between the 50% cardiomyocyte group and each other group. In these comparisons, we found a considerable effect of compositional changes on differential expression. Strikingly, a 10% reduction in the major cell type, a change smaller than that seen in cardiomyocyte abundance during MI in our data, caused 25% of transcripts to be identified as differentially expressed genes (DEGs) **(Figure 4C).** We repeated the analysis with different representations of proportions of the major cell type included as a term in the model, including unchanged percents, and both principal components and centered-log-ratio (CLR) transformations of cardiomyocyte proportions. In this updated model, nearly all gene expression changes are now associated with the terms representing cardiomyocyte abundance, rather than sample groups (which were blind to the underlying compositions of samples). These results demonstrate how comparisons of bulk gene expression can be skewed when sample groups have confounding variation in cell composition and how including compositional terms in DEG analysis enables more accurate attribution of expression changes.

Having validated a method of adjusting for compositional differences, we applied this analysis to our existing bulk RNAseq data. While each transformation of the compositional variable returned similarly low false positive rates in our simulation (**Supp. Fig. 2A**), we opted to include CLR-transformed proportions of cardiomyocytes and fibroblasts in the final DESeq2 model because CLRs conserve proper covariance structure of compositional data^64^, which could otherwise inflate associations when included as covariates in RNAseq analysis^65,66^. In this updated model we found that the cardiomyocyte abundance was associated with altered expression of hundreds of genes, although LCA ligation remained the lead variable. As before, LCA ligation was associated with the greatest number of DEGs (**Figure 4D**). Furthermore, inclusion of cell types in the model led to a broad reduction in overall significances of many of the genes previously attributed to the effect of LCA ligation or cmAKO during LCA ligation (compare **Figure 1D** to **Figure 4E**).

To better define how accounting for cellular abundance informs the interpretation of gene expression changes, we repeated the GO analyses using the results of the composition-adjusted DESeq2. By adjusting for these major cell types, some GO terms weakened or outright lost their associations with their original genotype or treatment variables (**Figure 4F, Supp. Figs. 2B-D**). For example, there was a reduction in the association found between MI and hallmark cardiomyocyte homeostatic processes, like cellular respiration (p = 2.30e-12 became p = 1.31e-03) and purine nucleotide metabolic process (p = 1.84e-4 became p = 0.0213) (**Figure 4G, Supp. Fig. 2C)**. Other GO terms saw more modest shifts, such as an increase in association of cmAKO X MI with biological processes indicative of cardiomyocyte state changes, including cardiac cell development (p = 1.83e-04 became p = 3.68e-05) and muscle cell differentiation (p = 1.83e-04 became p = 4.13e-05) (**Figure 4G, Supp. Fig. 2C**). Still other GO terms showed more considerable increases in the strength of their association with cmAKO x MI, such as processes related to coordinated heart function, such as heart contraction (p = 0.0897 to p = 3.68e-5), actin filament organization (p = 0.0658 to p = 6.32e-5), regulation of striated muscle contraction (p = 0.309 to p = 2.55e-4), potentially reflecting a reprogramming of CMs to a fetal-like state^67^. Together, these observations suggest that the cmAKO-specific response to MI involves a greater transcriptomic change in cardiomyocyte state when compared to WT mice.

Cell type abundances were significantly associated with many of the GO terms which decreased in their association with MI after correction for cell type. Specifically, alterations in fibroblast abundance were related to extracellular matrix-related processes like collagen fibril organization (p = 5.34e-03). Meanwhile, cardiomyocyte abundance was strongly associated with several processes related to energy metabolism, like cellular respiration (p = 2.13e-19) (**Figure 4H**). Additionally, we noticed an enrichment for cardiomyocyte association with terms indicative of inflammation, such as myeloid leukocyte migration (p = 4.81e-08) and adaptive immune response (p = 5.13e-07) (**Supplemental Data 3**). Reflecting the importance of cardiomyocytes as the major contributor to overall transcriptomic profile, the most significant GO terms were related to cardiomyocyte abundance (compare **Figure 4G** to **Figure 4H**, **Figure 4F-H**). These results indicate that the transcriptomic changes initially attributed to either LCA ligation alone or the unique response of cmAKO mice during LCA ligation may both be mediated by changes in cellular composition.

Taken together, our results suggest that our method was able to attribute transcriptomic changes due to deviation in cell type abundances from our compositional estimates but was not sufficiently powered to detect associations with cellular sub-states. We hypothesize that this may be due to the use of broad cell type annotations in our snRNAseq reference panel that lacks the resolution to correctly identify these more finely differentiated cellular populations.

### Spatial expression of cell type and cell state markers

To validate the ability of our model to predict regulation of transcripts within distinct cell types, we used RNAscope paired with immunohistochemistry. We designed probes for *Zbtb16* and *Pik3r1,* two highly regulated transcripts in the bulk RNAseq data that experienced large reductions in significance in their association with the cmAKO interaction with MI upon adjusting for shifts in cellular composition (**Figures 1D**, **4E**, **5A**). We then localized those probes within cardiomyocytes using immunofluorescent staining for sarcomeric alpha-actinin as an additional control for our methods.

**Figure 5:**
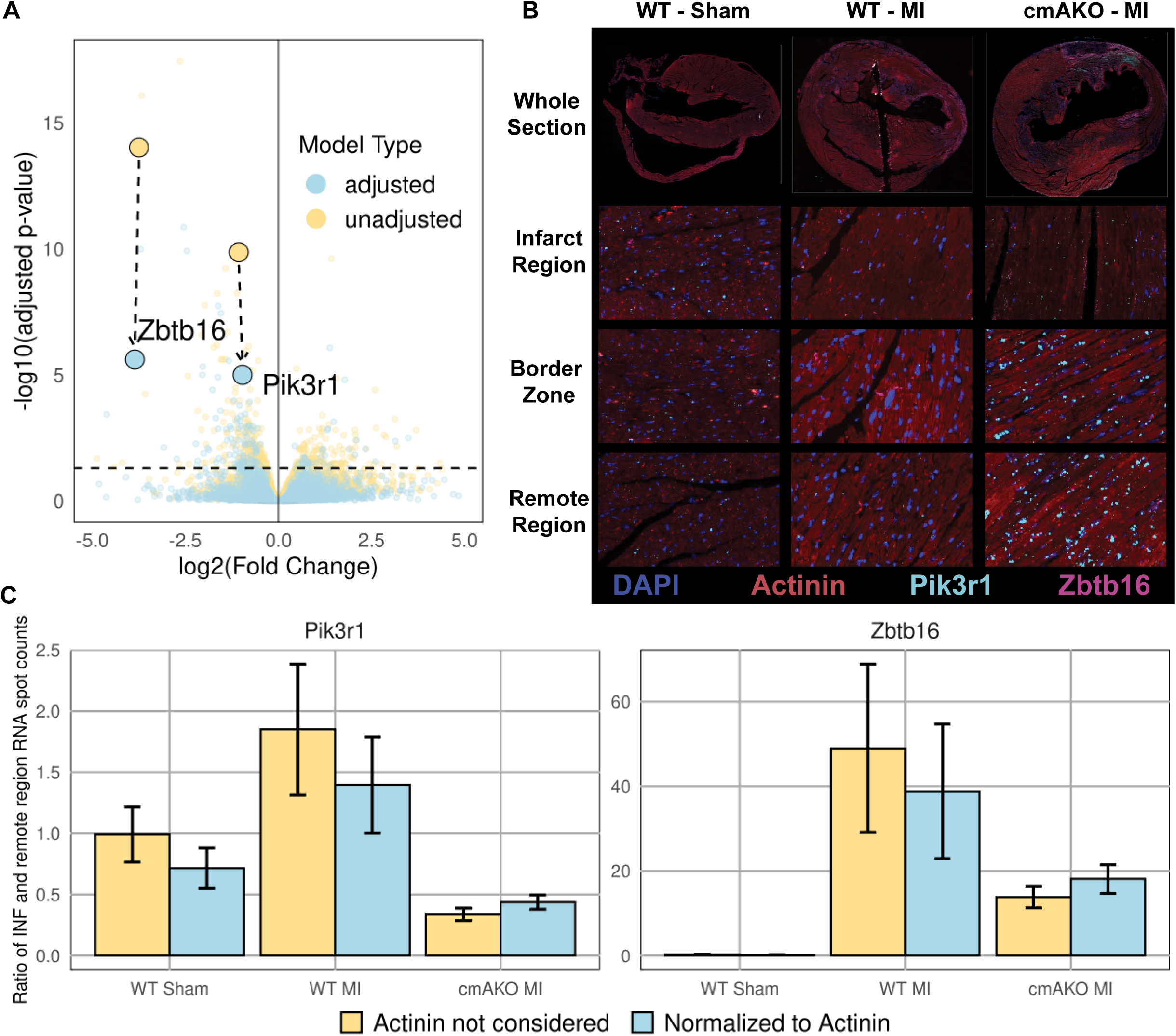
Fluorescent in-situ hybridization experimentally validates composition-aware differential gene expression results. **A)** Volcano plot showing expression changes associated with the unique response of cmAKO mice to myocardial infarction before (yellow) and after (blue) including terms for cardiomyocyte and fibroblast abundance in the DESeq2 model. Two highly significant genes, Zbtb16 and Pik3r1, are representative of a pattern of reduced significance in the updated model. **B)** RNAscope of Pik3r1 (cyan) and Zbtb16 (purple) overlayed with IHC showing DAPI (blue) and Actinin (red) in mouse left ventricular tissue. Representative images of the border zone, infract region, and a remote region from sham surgery control wild-type mice, wild-type mice after LCA ligation and cmAKO mice after LCA ligation. **C)** Adjusting for cardiomyocyte abundance minimizes differences in spatial expression of Pik3r1 and Zbtb16 in the infarct region, indicated by the post-adjustment expression increases in cmAKO MI and reductions in WT MI. Five representative regions were evaluated in each zone from panel B and the the number of expression spots were tallied for each gene. Actinin was used as a proxy for cardiomyocyte abundance and spot counts were normalized to actinin abundance (blue) or not (yellow). To control for inter-slide technical variation, spot counts were normalized to slide-matched measures in a remote region (shown on the y-axis).

Our predictive model indicated that *Zbtb16* (a zinc finger transcription factor), and *Pik3r1* (PI 3-kinase regulatory subunit 1) were regulated in a genotype-specific manner within cardiomyocytes after MI. Both transcripts are known to be expressed in cardiomyocytes^53,68^, but no previous studies have identified them as downstream targets of α1-ARs. We found that both *Zbtb16* and *Pik3r1* were expressed in cardiomyocytes under all conditions. Control hearts (**Figure 5B**) displayed uniform transcript expression across tissue regions. Consistent with our predictions, transcript abundance of cardiomyocyte *Zbtp16* was relatively low in ligated WT mouse hearts compared to the robust upregulation in hearts from ligated cmAKO mice. The pattern of *Pik3r1* expression was identical, although the magnitude of differential expression was qualitatively lower, also consistent with our predictions.

To quantify these effects, we evaluated five representative subregions within the border zone, infarct region, and remote region (or matched locations) of each slide. For each subregion, we tallied the number of expression spots of each gene, as well as the area occupied by actinin, which was used as a proxy for total cardiomyocyte cross-sectional area. To account for differences in cardiomyocyte abundance (**Supp. Fig. 3**), we then normalized expression spot counts to the area occupied by actinin in each subregion. Due to the technical variation present between each image, we further normalized these values to those from the remote region of each sample. We then observed that accounting for cardiomyocyte abundance reduced the differences in spatial expression both *Zbtb16* and *Pik3r1* between cmAKO and WT mice during myocardial infarction (**Figure 5C**). This finding is consistent with the predictions made in the composition-aware differential expression analysis, which suggested cell type proportion shifts were driving the altered expression of *Zbtb16* and *Pik3r1*.

## Discussion

Changes in cardiac cell composition that occur during cardiac remodeling are a hallmark of cardiac dysfunction. Notably, loss of cardiomyocytes and proliferation of fibroblasts has been previously reported in myocardial infarction, trans-aortic constriction and beta-adrenergic overdrive models of heart disease^69,70,71^. Quantitative analyses of the degree of remodeling in the heart remains complicated due to a variety of technical and biological factors. In this study, we have developed a computational method that accounts for transcriptomic changes in bulk RNA sequencing data by delineating cell-type specific changes. We explore the efficacy of this approach in bulk RNAseq datasets drawn from wild type and cardiomyocyte-specific α1A-AR knockout mice (CM-α1A-KO) subjected to myocardial infarction or sham surgery.

A growing body of literature demonstrates that α1A-ARs play adaptive and protective roles in cell culture and animal models^2,72^. We recently found that selective antagonists of α1A-ARs such as tamsulosin (Flomax), used commonly to treat lower urinary tract symptoms related to benign prostatic hypertrophy, are associated with a small but statistically significant increase in one-year mortality in a subset of the Medicare database^73^. To understand the primacy of cardiomyocyte α1A-ARs in these findings, we created a mouse line (cmAKO) with cardiomyocyte-specific deletion of the α1A-AR and found that cmAKO mice had markedly increased mortality and exacerbated pathological ventricular remodeling following myocardial infarction (MI)^9^.

Applying our computational deconvolution algorithm to this model enabled us to pinpoint how shifts in cell populations — like increases in fibroblasts and decreases in cardiomyocytes — directly influence gene expression changes during cardiac injury. Our results produce novel insights into the specific cellular mechanisms affected by α1-AR and provide an invaluable toolkit for similar transcriptomic studies, offering a robust, scalable strategy to interpret complex biological data when there are shifts in cell type heterogeneity.

Our method introduces a significant improvement in the analysis of bulk RNA sequencing data by accurately estimating the contributions of distinct cell types within heterogeneous cardiac samples. This approach addresses a common limitation in traditional bulk RNAseq analysis, where gene expression differences due to changes in cell state are difficult to distinguish from those due to changes in cellular abundance^10^. As cellular remodeling is an expected feature of many cardiac diseases, transcriptomic studies in the heart often identify cell-type specific markers as significantly differentially expressed. These changes in expression are frequently attributed to changes in biological activity rather than relative cellular abundance^74,75^. When cellular makeup has been addressed in cardiac bulk RNAseq studies, changes in cellular abundance have often been post-hoc inferred through alterations in cell-type-specific transcripts^72,75^. By contrast, our approach aims to precisely model the entire cellular state of the heart before comparisons of gene expression are performed. In our approach, distinguishing these contributions represents an advancement to the field that offers a straightforward means of interpreting complex transcriptomic data obtained from bulk tissue, allowing researchers to quantify the contributions of individual cell types to total gene expression.

Researchers have produced a tremendous amount of bulk RNAseq data in the last decade and it remains a first-line tool for profiling transcriptomes. More than 11,000 bulk RNA-seq datasets have been deposited in NCBI’s Gene Expression Omnibus (GEO) and 79% of NGS data deposited in 2022 was from bulk RNA datasets, rather than single-cell studies ^76^. As such, this approach opens new avenues for both retrospective and prospective research projects. Existing cohorts may be reanalyzed, potentially uncovering novel insights that were previously obscured due to changes in cellular abundance. Furthermore, by correcting for cell-type abundances, our method helps to refine models of disease mechanisms by delineating effects to their cell-type of origin, thereby improving the predictive accuracy of transcriptomic markers of disease.

While our computational method significantly enhances the analysis of bulk RNA sequencing data from heterogeneous samples, it is not without limitations. One key challenge is the reliance on accurate cell-type-specific markers. Incorrect or suboptimal marker selection can lead to inaccurate deconvolution results, which may obscure true cellular contributions to gene expression changes. Ensuring the precision of these markers requires extensive validation, which can be time-intensive and is limited by the availability of high-quality reference datasets. Indeed, this method is beholden to both the cell types and states present in the supplied reference, which may require researchers to generate their own references if applied in more niche biological contexts. If cell types with expected proportion changes are excluded from the study, it is likely that their effects will be misattributed to other, highly correlated cell types, as seen in the case of immune-cell-specific mechanisms that are inappropriately associated with cardiomyocytes in our analysis. Moreover, while our method improves the resolution of cellular contributions, the broader field of cell type deconvolution struggles to maintain accuracy as the number of included cell types and/or their similarity to one another increases^74^.

In this study, we defined the transcriptomic signatures that result from the absence of cardiomyocyte α1A-ARs in the uninjured and post-infarct state. Leveraging a novel computational approach, we accurately estimated cell type-specific contributions within bulk RNA sequencing data. Further, we validated the method by visualizing the spatial expression of two representative transcripts, recapitulating the findings from our computational model. This method enabled us to precisely dissect how shifts in cellular composition, such as increases in fibroblasts and decreases in cardiomyocytes, directly impact gene expression during cardiac stress. The resulting composition-aware dataset identified novel associations with pathways and individual transcripts that will informfuture mechanistic studies to expand our understanding of the cardioprotective effects of 1A-ARs. Additionally, these results pose implications for how clinicians may consider the tissue-level effects of therapeutics mediated by adrenergic receptors. Future efforts to define cellular composition in cardiac tissue may benefit from benchmarking the accuracy of deconvolution algorithms when applied to a broader range of cellular states and types, as prior studies have found current methods to vary in their performance across different tissue contexts^77,78^. This study not only contributes to our understanding of the molecular dynamics within the heart but also offers a robust, scalable strategy to uncover hidden insights in pre-existing and future transcriptomic data.

## Acknowledgements

We acknowledge the grants that supported this research: BG (T32HL069768), CDR (R01HL162636, R00HL138301), TMV NIH grants HL105699 and HL 159086, BCJ (2R01 HL140067). We thank Rachel Sharp and Sarah Lester for their constructive comments on this manuscript, Michael Love, Ph.D. for his helpful advice regarding statistical approaches, and the Roy J Carver Biotechnology Center, and the Technology Center for Genomics and Bioinformatics for their help with performing the RNA sequencing. We acknowledge Biorender for their support in generating figures 1-4.

