## Supplementary Figures for "Novel Insights into Post-Myocardial Infarction Cardiac Remodeling through Algorithmic Detection of Cell-Type Composition Shifts"

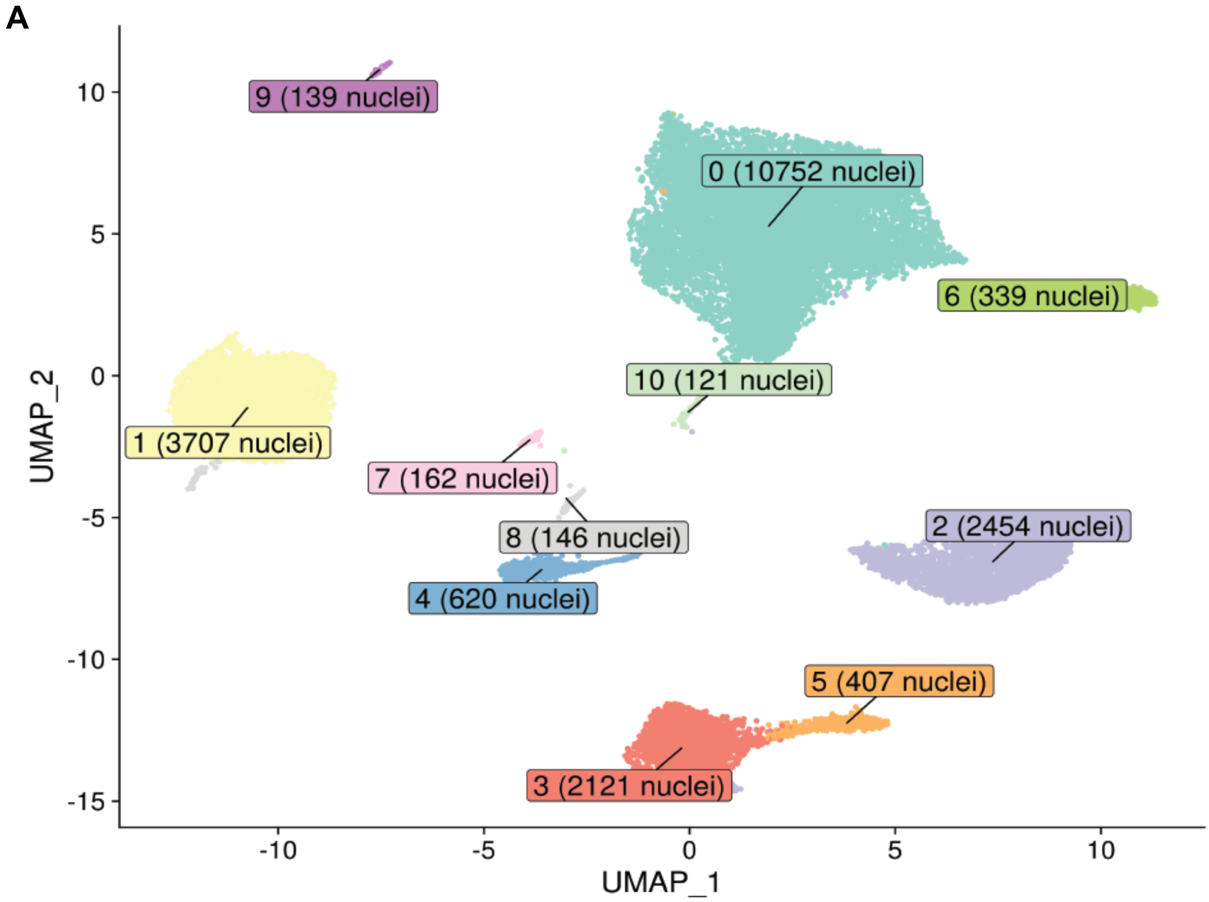


**Supplementary Figure 1:** UMAP of single nucleus RNA sequencing dataset after intial quality control involving doublet detection, ambient RNA removal, filtering by features such as gene counts and mitocondrial transcript abundance. Clusters 6 - 10 were excluded from the final analysis due to their low nuclei counts and poor annotation to known cell types. Clusters 3 and 5 were merged due to their similar marker profiles.

**Supplementary Figure 2:** Three modalities of cell type proportions were considered for use in differential gene expression testing: center log ratio (CLR) transformation, prinicpal components (PCs) from principal component analysis, and untransformed proportions. For the CLR and untransformed approaches, the cardiomyocyte-asssociated value was included as a variable in the DESeq2 model, while the first PC was used for the PC-based approach. See methods and Figure 4A for more information.

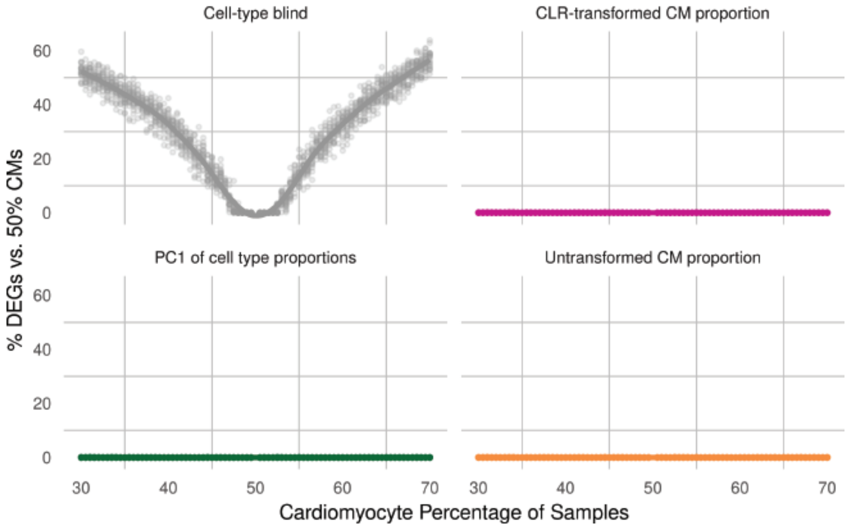

**Supplementary Figure 3: Composition-aware gene ontology analysis of α1A-AR knockout mice during myocardial infarction. A** and **B,** GO terms associated with myocardial infarction and cmAKOxMI ordered by largest reduction (**A**) and gain (**B**) in significance after accounting for cardiomyocyte and fibroblast abundance. **C** and **D,** Extended listing of top GO terms post-adjustment showing significant associations with myocardial infarction and cmAKOxMI (**C**), and with changes in cardiomyocytes and fibroblasts (**D**).

**A**

GO terms associated with WT and cmAKO responses to MI

By significance lost after cell-type-adjustment

| | Cell-type not considered | Cell-type adjusted | |
|---|---|---|---|
| | p-value (Q-score) | p-value (Q-score) | ΔQ-score |
| **Myocardial Infarction** | | | |
| Aerobic Respiration | 8.77e-13 (12.06) | 8.63e-04 (3.0640) | -8.99 |
| Oxidative Phosphorylation | 1.26e-14 (13.90) | 8.96e-06 (5.0478) | -8.85 |
| Cellular Respiration | 2.30e-12 (11.64) | 1.31e-03 (2.8837) | -8.75 |
| Purine Nucleotide Metabolic Process | 1.84e-10 (9.74) | 2.13e-02 (1.6723) | -8.06 |
| Energy Derivation By Oxidation Of Organic Compounds | 2.80e-11 (10.55) | 2.50e-03 (2.6024) | -7.95 |
| Purine-Containing Compound Metabolic Process | 6.55e-10 (9.18) | 4.59e-02 (1.3381) | -7.85 |
| Atp Metabolic Process | 1.81e-11 (10.74) | 9.33e-04 (3.0302) | -7.71 |
| Purine Ribonucleotide Metabolic Process | 1.91e-10 (9.72) | 9.63e-03 (2.0165) | -7.70 |
| Purine Nucleoside Triphosphate Metabolic Process | 9.30e-11 (10.03) | 3.21e-03 (2.4938) | -7.54 |
| Nucleotide Metabolic Process | 1.65e-09 (8.78) | 3.43e-02 (1.4647) | -7.32 |
| Ribonucleotide Metabolic Process | 9.37e-10 (9.03) | 1.93e-02 (1.7137) | -7.31 |
| Ribose Phosphate Metabolic Process | 2.19e-09 (8.66) | 3.22e-02 (1.4925) | -7.17 |
| Nucleoside Phosphate Metabolic Process | 2.77e-09 (8.56) | 3.70e-02 (1.4320) | -7.13 |
| Purine-Containing Compound Biosynthetic Process | 5.68e-12 (11.25) | 6.87e-05 (4.1632) | -7.08 |
| Atp Synthesis Coupled Electron Transport | 4.57e-10 (9.34) | 4.77e-03 (2.3212) | -7.02 |
| Mitochondrial Atp Synthesis Coupled Electron Transport | 4.57e-10 (9.34) | 4.77e-03 (2.3212) | -7.02 |
| Nucleoside Triphosphate Metabolic Process | 3.34e-10 (9.48) | 3.24e-03 (2.4896) | -6.99 |
| Purine Ribonucleoside Triphosphate Metabolic Process | 1.91e-10 (9.72) | 1.54e-03 (2.8132) | -6.91 |
| Atp Biosynthetic Process | 2.80e-11 (10.55) | 2.21e-04 (3.6557) | -6.90 |
| Respiratory Electron Transport Chain | 6.55e-10 (9.18) | 3.33e-03 (2.4782) | -6.71 |
| **cmAKO:MI** | | | |
| Regulation Of Small Gtpase Mediated Signal Transduction | 1.26e-03 (2.90) | 4.24e-01 (0.3728) | -2.53 |
| Regulation Of Ras Protein Signal Transduction | 1.35e-03 (2.87) | 4.40e-01 (0.3568) | -2.51 |
| Positive Regulation Of Ras Protein Signal Transduction | 1.70e-03 (2.77) | 5.10e-01 (0.2928) | -2.48 |
| Regulation Of Leukocyte Mediated Immunity | 3.63e-03 (2.44) | 9.17e-01 (0.0374) | -2.40 |
| Cell Killing | 2.61e-03 (2.58) | 5.70e-01 (0.2440) | -2.34 |
| Myeloid Leukocyte Mediated Immunity | 4.46e-03 (2.35) | 9.29e-01 (0.0320) | -2.32 |
| Positive Regulation Of Cell Activation | 2.39e-03 (2.62) | 4.40e-01 (0.3561) | -2.27 |
| Leukocyte Mediated Immunity | 5.09e-03 (2.29) | 9.24e-01 (0.0344) | -2.26 |
| Positive Regulation Of Small Gtpase Mediated Signal Transduction | 3.11e-03 (2.51) | 5.35e-01 (0.2714) | -2.24 |
| Interleukin-1 Production | 6.76e-04 (3.17) | 1.11e-01 (0.9564) | -2.21 |
| Regulation Of Tumor Necrosis Factor Production | 1.71e-03 (2.77) | 2.64e-01 (0.5780) | -2.19 |
| Regulation Of Tumor Necrosis Factor Superfamily Cytokine Production | 1.95e-03 (2.71) | 2.69e-01 (0.5696) | -2.14 |
| Negative Regulation Of Mapk Cascade | 2.61e-03 (2.58) | 3.51e-01 (0.4551) | -2.13 |
| Leukocyte Migration | 1.26e-03 (2.90) | 1.63e-01 (0.7867) | -2.11 |
| Interleukin-1 Beta Production | 5.02e-04 (3.30) | 6.32e-02 (1.1992) | -2.10 |
| Tumor Necrosis Factor Production | 2.52e-03 (2.60) | 2.84e-01 (0.5466) | -2.05 |
| Tumor Necrosis Factor Superfamily Cytokine Production | 2.61e-03 (2.58) | 2.93e-01 (0.5326) | -2.05 |
| Positive Regulation Of Glucose Metabolic Process | 3.51e-03 (2.45) | 3.55e-01 (0.4494) | -2.01 |
| Myeloid Cell Activation Involved In Immune Response | 7.73e-03 (2.11) | 7.23e-01 (0.1409) | -1.97 |
| Regulation Of T Cell Cytokine Production | 5.74e-03 (2.24) | 4.63e-01 (0.3347) | -1.91 |

**B**

Most significantly increased terms after adjustment

| | Cell-type not considered | Cell-type adjusted | |
|---|---|---|---|
| | p-value (Q-score) | p-value (Q-score) | ΔQ-score |
| **Myocardial Infarction** | | | |
| Positive Regulation Of Leukocyte Migration | 8.07e-05 (4.093) | 1.55e-07 (6.81) | 2.72 |
| Leukocyte Chemotaxis | 8.11e-06 (5.091) | 2.82e-08 (7.55) | 2.46 |
| Myeloid Leukocyte Migration | 9.93e-09 (8.003) | 3.46e-11 (10.46) | 2.46 |
| Humoral Immune Response | 1.25e-09 (8.902) | 9.59e-12 (11.02) | 2.12 |
| Cell Chemotaxis | 1.27e-06 (5.896) | 1.06e-08 (7.98) | 2.08 |
| Leukocyte Migration | 1.05e-09 (8.977) | 9.59e-12 (11.02) | 2.04 |
| Granulocyte Chemotaxis | 3.37e-07 (6.473) | 3.68e-09 (8.43) | 1.96 |
| Regulation Of Bmp Signaling Pathway | 7.01e-02 (1.154) | 1.36e-03 (2.87) | 1.71 |
| Complement Activation | 1.48e-09 (8.828) | 3.81e-11 (10.42) | 1.59 |
| Regulation Of Leukocyte Migration | 1.38e-05 (4.859) | 3.55e-07 (6.45) | 1.59 |
| Taxis | 1.16e-07 (6.936) | 2.98e-09 (8.53) | 1.59 |
| Chemotaxis | 9.45e-08 (7.025) | 2.55e-09 (8.59) | 1.57 |
| Peptide Cross-Linking | 1.55e-03 (2.810) | 5.67e-05 (4.25) | 1.44 |
| Fibroblast Proliferation | 2.47e-03 (2.607) | 1.07e-04 (3.97) | 1.36 |
| Mononuclear Cell Migration | 5.19e-03 (2.285) | 2.33e-04 (3.63) | 1.35 |
| Cell Recognition | 1.02e-01 (0.990) | 4.60e-03 (2.34) | 1.35 |
| Metaphase Chromosome Alignment | 8.47e-02 (1.072) | 4.83e-03 (2.32) | 1.24 |
| Gland Morphogenesis | 2.10e-02 (1.677) | 1.26e-03 (2.90) | 1.22 |
| Prostanoid Metabolic Process | 4.33e-05 (4.363) | 2.78e-06 (5.56) | 1.19 |
| Modulation Of Process Of Another Organism | 1.45e-02 (1.840) | 9.41e-04 (3.03) | 1.19 |
| **cmAKO:MI** | | | |
| Heart Contraction | 8.97e-02 (1.047) | 3.68e-05 (4.43) | 3.39 |
| Heart Process | 7.67e-02 (1.115) | 6.32e-05 (4.20) | 3.08 |
| Regulation Of Striated Muscle Contraction | 3.09e-01 (0.510) | 2.55e-04 (3.59) | 3.08 |
| Actin Filament Organization | 6.58e-02 (1.182) | 6.32e-05 (4.20) | 3.02 |
| Regulation Of Cardiac Muscle Contraction | 2.16e-01 (0.665) | 2.55e-04 (3.59) | 2.93 |
| Regulation Of Heart Contraction | 5.57e-02 (1.254) | 7.05e-05 (4.15) | 2.90 |
| Import Across Plasma Membrane | 2.99e-01 (0.525) | 6.96e-04 (3.16) | 2.63 |
| Cardiac Muscle Contraction | 2.10e-01 (0.677) | 5.68e-04 (3.25) | 2.57 |
| Striated Muscle Contraction | 2.46e-01 (0.609) | 1.22e-03 (2.91) | 2.31 |
| Developmental Cell Growth | 8.92e-02 (1.050) | 5.68e-04 (3.25) | 2.20 |
| Regulation Of Muscle Contraction | 1.66e-01 (0.781) | 1.34e-03 (2.87) | 2.09 |
| Muscle Tissue Development | 4.46e-03 (2.351) | 3.68e-05 (4.43) | 2.08 |
| Inorganic Cation Import Across Plasma Membrane | 4.49e-01 (0.348) | 4.13e-03 (2.38) | 2.04 |
| Inorganic Ion Import Across Plasma Membrane | 4.49e-01 (0.348) | 4.13e-03 (2.38) | 2.04 |
| Regulation Of Actin Filament-Based Process | 1.11e-01 (0.956) | 1.52e-03 (2.82) | 1.86 |
| Actin Filament-Based Movement | 2.63e-01 (0.581) | 3.74e-03 (2.43) | 1.85 |
| Negative Regulation Of Locomotion | 1.66e-01 (0.781) | 2.64e-03 (2.58) | 1.80 |
| Fat Cell Differentiation | 6.20e-01 (0.207) | 1.50e-02 (1.82) | 1.62 |
| Axon Extension Involved In Axon Guidance | 3.39e-01 (0.470) | 9.16e-03 (2.04) | 1.57 |
| Neuron Projection Extension Involved In Neuron Projection Guidance | 3.39e-01 (0.470) | 9.16e-03 (2.04) | 1.57 |

**C**

Cell-type adjusted GO terms associated with WT and cmAKO responses to MI

| | Cell-type not considered | Cell-type adjusted | |
|---|---|---|---|
| | p-value (Q-score) | p-value (Q-score) | ΔQ-score |
| **Myocardial Infarction** | | | |
| Extracellular Matrix Organization | 1.25e-19 (18.903) | 1.02e-17 (16.99) | -1.9104 |
| Extracellular Structure Organization | 1.25e-19 (18.903) | 1.02e-17 (16.99) | -1.9104 |
| External Encapsulating Structure Organization | 1.25e-19 (18.903) | 1.02e-17 (16.99) | -1.9104 |
| Humoral Immune Response | 1.25e-09 (8.902) | 9.59e-12 (11.02) | 2.1161 |
| Leukocyte Migration | 1.05e-09 (8.977) | 9.59e-12 (11.02) | 2.0407 |
| Neutrophil Migration | 3.57e-11 (10.447) | 2.67e-11 (10.57) | 0.1258 |
| Response To Bacterium | 1.10e-13 (12.958) | 2.67e-11 (10.57) | -2.3856 |
| Myeloid Leukocyte Migration | 9.93e-09 (8.003) | 3.46e-11 (10.46) | 2.4585 |
| Complement Activation | 1.48e-09 (8.828) | 3.81e-11 (10.42) | 1.5912 |
| Granulocyte Migration | 6.55e-10 (9.184) | 4.90e-11 (10.31) | 1.1256 |
| Adaptive Immune Response | 1.31e-10 (9.882) | 9.92e-11 (10.00) | 0.1213 |
| Leukocyte Mediated Immunity | 3.12e-10 (9.506) | 3.74e-10 (9.43) | -0.0784 |
| Neutrophil Chemotaxis | 1.16e-08 (7.937) | 9.62e-10 (9.02) | 1.0798 |
| Myeloid Leukocyte Activation | 1.01e-09 (8.995) | 1.56e-09 (8.81) | -0.1890 |
| Chemotaxis | 9.45e-08 (7.025) | 2.55e-09 (8.59) | 1.5686 |
| Taxis | 1.16e-07 (6.936) | 2.98e-09 (8.53) | 1.5906 |
| Granulocyte Chemotaxis | 3.37e-07 (6.473) | 3.68e-09 (8.43) | 1.9619 |
| Cell Chemotaxis | 1.27e-06 (5.896) | 1.06e-08 (7.98) | 2.0794 |
| Adaptive Immune Response Based On Somatic Recombination Of Immune Receptors Built From Immunoglobulin Superfamily Domains | 4.56e-09 (8.341) | 1.06e-08 (7.98) | -0.3657 |
| Cell Activation Involved In Immune Response | 1.16e-08 (7.937) | 1.32e-08 (7.88) | -0.0584 |
| **cmAKO:MI** | | | |
| Cardiac Cell Development | 1.83e-04 (3.737) | 3.68e-05 (4.43) | 0.6975 |
| Cardiac Muscle Cell Development | 1.83e-04 (3.737) | 3.68e-05 (4.43) | 0.6975 |
| Muscle Tissue Development | 4.46e-03 (2.351) | 3.68e-05 (4.43) | 2.0837 |
| Muscle Cell Development | 1.26e-03 (2.900) | 3.68e-05 (4.43) | 1.5348 |
| Heart Contraction | 8.97e-02 (1.047) | 3.68e-05 (4.43) | 3.3875 |
| Cardiac Muscle Cell Differentiation | 1.83e-04 (3.737) | 4.13e-05 (4.38) | 0.6472 |
| Muscle Cell Differentiation | 1.83e-04 (3.737) | 4.13e-05 (4.38) | 0.6472 |
| Heart Process | 7.67e-02 (1.115) | 6.32e-05 (4.20) | 3.0847 |
| Actin Filament Organization | 6.58e-02 (1.182) | 6.32e-05 (4.20) | 3.0178 |
| Striated Muscle Tissue Development | 5.02e-04 (3.299) | 6.32e-05 (4.20) | 0.9004 |
| Cardiac Muscle Tissue Development | 5.02e-04 (3.299) | 6.84e-05 (4.16) | 0.8656 |
| Regulation Of Heart Contraction | 5.57e-02 (1.254) | 7.05e-05 (4.15) | 2.8978 |
| Striated Muscle Cell Differentiation | 5.02e-04 (3.299) | 9.23e-05 (4.03) | 0.7357 |
| Cardiocyte Differentiation | 3.10e-04 (3.508) | 9.33e-05 (4.03) | 0.5221 |
| Regulation Of Cardiac Muscle Contraction | 2.16e-01 (0.665) | 2.55e-04 (3.59) | 2.9280 |
| Regulation Of Striated Muscle Contraction | 3.09e-01 (0.510) | 2.55e-04 (3.59) | 3.0827 |
| Muscle System Process | 5.02e-04 (3.299) | 4.32e-04 (3.36) | 0.0656 |
| Regulation Of Heart Rate | 7.20e-03 (2.143) | 5.68e-04 (3.25) | 1.1031 |
| Cardiac Muscle Contraction | 2.10e-01 (0.677) | 5.68e-04 (3.25) | 2.5683 |
| Developmental Cell Growth | 8.92e-02 (1.050) | 5.68e-04 (3.25) | 2.1960 |

**D**

GO terms associated with cardiomyocytes and fibroblasts

| | p-value (Q-score) |
|---|---|
| **Cardiomyocytes** | |
| Cellular Respiration | 2.13e-19 (18.67) |
| Purine Nucleotide Metabolic Process | 5.87e-18 (17.23) |
| Purine-Containing Compound Metabolic Process | 7.56e-18 (17.12) |
| Nucleotide Metabolic Process | 7.56e-18 (17.12) |
| Energy Derivation By Oxidation Of Organic Compounds | 1.45e-17 (16.84) |
| Nucleoside Phosphate Metabolic Process | 1.45e-17 (16.84) |
| Aerobic Respiration | 2.79e-17 (16.55) |
| Generation Of Precursor Metabolites And Energy | 9.13e-16 (15.04) |
| Nucleobase-Containing Small Molecule Metabolic Process | 6.35e-15 (14.20) |
| Tricarboxylic Acid Cycle | 1.83e-14 (13.74) |
| Purine Ribonucleotide Metabolic Process | 4.88e-14 (13.31) |
| Ribonucleotide Metabolic Process | 7.40e-14 (13.13) |
| Respiratory Electron Transport Chain | 7.40e-14 (13.13) |
| Ribose Phosphate Metabolic Process | 2.40e-13 (12.62) |
| Electron Transport Chain | 3.25e-13 (12.49) |
| Monocarboxylic Acid Metabolic Process | 9.56e-12 (11.02) |
| Fatty Acid Beta-Oxidation | 1.47e-11 (10.83) |
| Fatty Acid Catabolic Process | 1.47e-11 (10.83) |
| Atp Synthesis Coupled Electron Transport | 2.05e-11 (10.69) |
| Mitochondrial Atp Synthesis Coupled Electron Transport | 2.05e-11 (10.69) |
| **Fibroblasts** | |
| Collagen Fibril Organization | 5.34e-03 (2.27) |
| Extracellular Matrix Organization | 1.11e-02 (1.96) |
| Extracellular Structure Organization | 1.11e-02 (1.96) |
| External Encapsulating Structure Organization | 1.11e-02 (1.96) |
| Epithelial Cell Proliferation | 1.39e-02 (1.86) |
| Skeletal System Development | 1.65e-02 (1.78) |
| Endoderm Formation | 1.99e-02 (1.70) |
| Glial Cell Migration | 2.74e-02 (1.56) |
| Muscle Contraction | 2.80e-02 (1.55) |
| Pyrimidine Nucleoside Triphosphate Biosynthetic Process | 2.80e-02 (1.55) |
| Kidney Development | 2.80e-02 (1.55) |
| Myeloid Leukocyte Differentiation | 3.34e-02 (1.48) |
| Renal System Development | 3.34e-02 (1.48) |
| Deoxyribonucleotide Biosynthetic Process | 3.34e-02 (1.48) |
| 2'-Deoxyribonucleotide Biosynthetic Process | 3.34e-02 (1.48) |
| Deoxyribose Phosphate Biosynthetic Process | 3.34e-02 (1.48) |
| Pyrimidine Nucleoside Triphosphate Metabolic Process | 4.44e-02 (1.35) |

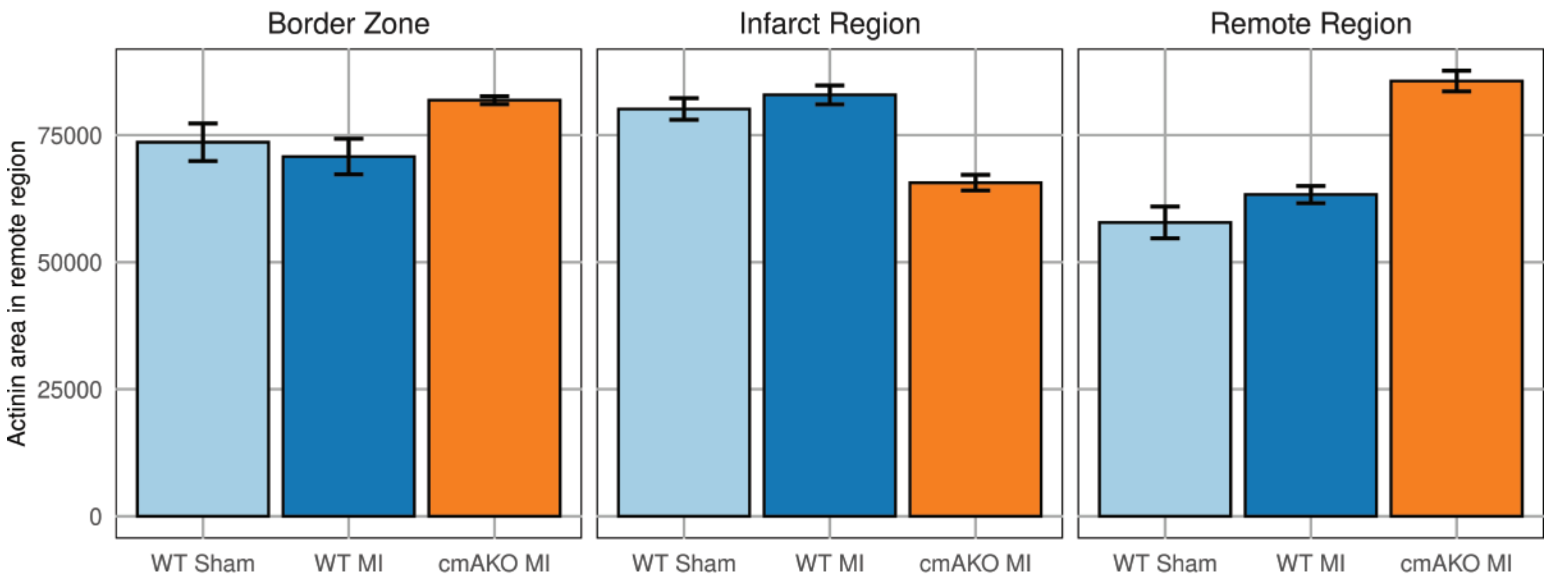


**Supplementary Figure 4:** Area occupied by sarcomeric actinin in immunohistological staining of cardiac cross sections. Five representative regions were evaluated in each zone from Figure 5B and modest variation is seen within each sample between regions. Area was measured in pixel counts.
